# Time-dependent inhibition of covert shift of attention

**DOI:** 10.1101/2020.07.17.207928

**Authors:** Antimo Buonocore, Niklas Dietze, Robert D. McIntosh

**Author notes:** Corresponding author: Antimo Buonocore, Werner Reichardt Centre for Integrative Neuroscience, Otfried-Müller Str. 25, Tübingen, 72076, Germany. equal contribution (Antimo Buonocore, Niklas Dietze).

## Abstract

Visual transients can interrupt overt orienting by abolishing the execution of a planned eye movement due about 90 ms later, a phenomenon known as saccadic inhibition (SI). It is not known if the same inhibitory process might influence covert orienting in the absence of saccades, and consequently alter visual perception. In Experiment 1 (n=14), we measured orientation discrimination during a covert orienting task in which an uninformative exogenous visual cue preceded the onset of an oriented probe by 140 to 290 ms. In half of the trials the onset of the probe was accompanied by a brief irrelevant flash, a visual transient that would normally induce SI. We report a time-dependent inhibition of covert orienting in which the irrelevant flash impaired orientation discrimination accuracy when the probe followed the cue by 190 and 240 ms. The interference was more pronounced when the cue was incongruent with the probe location, suggesting an impact on the reorienting component of the attentional shift. In Experiment 2 (n=12), we tested whether the inhibitory effect of the flash could occur within an earlier time range, or only within the later, reorienting range. We presented probes at congruent cue locations in a time window between 50 to 200 ms. Similarly to Experiment 1, discrimination performance was altered at 200 ms after the cue. We suggest that covert attention may be susceptible to similar inhibitory mechanisms that generate SI, especially in later stages of attentional shifting (> 200ms after a cue), typically associated with reorienting.

## Introduction

The primary function of visual processing is to guide efficient interactions with the surrounding world. This requires us to rapidly integrate new sensory information with ongoing motor behaviour to generate appropriate responses such as inhibiting a planned action in order to orient to a novel stimulus. One of the most striking examples of the fast integration of visual information into a motor command is saccadic inhibition (SI) (Bompas and Sumner 2011; Buonocore and McIntosh 2008; Buonocore and McIntosh 2012; Edelman and Xu 2009; Reingold and Stampe 1999; Reingold and Stampe 2002). It has been consistently observed that visual transient events interrupt ongoing eye movement behaviour, such that saccades that would otherwise be launched about 90 ms later are inhibited, or delayed. This is visualized as a distinct dip in saccadic frequency around 90 ms after the onset of the visual event, and followed by a rebound period in which the probability of making an eye movement is increased (Reingold and Stampe 2002). From a neuronal perspective, SI might be the result of an interruption in the build-up activity of superior colliculus (SC) neurons (Buonocore et al. 2021; Dorris et al. 2007; Munoz and Istvan 1998) coding for the location in space to which the saccade is planned (Munoz and Guitton 1991; Munoz and Wurtz 1993; Munoz and Wurtz 1995).

The exact mechanisms behind SI are still unknown, but there are strong hypotheses suggesting the involvement of omnipause neurons (OPN) in the nucleus raphe interpositus (Büttner-Ennever et al. 1988). OPNs are a small group of neurons that fire tonically during fixation and pause their activity abruptly just before and during saccades (Cohen and Henn 1972; Keller 1974; Luschei and Fuchs 1972). Sudden reactivation of OPNs following the onset of a visual transient might send an inhibitory signal interfering with the build-up activity in upstream oculomotor areas (Buonocore et al. 2020). Despite being one of the most reliable phenomena in oculomotor behaviour, the ecological function of SI is not fully understood.

We recently asked whether this interruption of ongoing oculomotor behaviour might have functional benefits in terms of facilitating overt reorienting to a new location in space (Buonocore et al. 2017c). For this purpose, we adapted the well-known double-step paradigm (Becker and Jürgens 1979; Lisberger et al. 1975), asking participants to make an eye movement in response to a sudden-onset visual target, which sometimes jumped to a new location before the first saccade could be launched. In half of the trials the jump was accompanied by the presentation of a brief (30 ms) visual flash at the top and bottom of the screen. These “flash-jump” trials induced strong saccadic inhibition, and led to a higher rate of successful reorienting to the new target location. We suggested that SI allowed the oculomotor system the time for a decisional process to change the planned response. Following this empirical demonstration, new models of saccadic inhibition have been proposed, aiming to unify inhibition with the response inhibition observed in countermanding tasks (Bompas et al. 2020; Salinas and Stanford 2018). These findings and models have advanced our understanding of the oculomotor system by suggesting the existence of a common inhibitory signal, driven by new visual onsets, capable of interrupting ongoing orienting. Nonetheless, they have been concerned exclusively with overt orienting (saccades), leaving open the question of whether covert orienting could be similarly affected.

Moving the eyes is not the only way to modulate sampling of visual information from the surroundings. Numerous studies have shown perceptual benefits at locations that have been previously cued, even when no eye movement is made, confirming that it is possible to orient attention covertly (Carrasco 2011; Posner 1980; see: Posner 2015 for a review). These studies have mostly shown speeding of simple detection responses, but covert orienting can also improve the discimination of spatial frequency and lower the contrast threshold for orientation discimination (Barbot et al. 2012; Cameron et al. 2002; Carrasco 2011; Fernández et al. 2019; Lee et al. 1999; Pestilli and Carrasco 2005; Solomon 2004). These changes in visual sensitivity are tightly coupled with the planning of eye movements (Buonocore et al. 2017b; Deubel and Schneider 1996; Hoffman and Subramaniam 1995; Remington 1980; Shepherd et al. 1986; Zhao et al. 2012) as implied by the observation of pre-saccadic perceptual enhancements at the target location of an upcoming saccade (Hanning et al. 2019; Kowler et al. 1995; Li et al. 2016; Li et al. 2019; Montagnini and Castet 2007; Ohl et al. 2017; Rolfs and Carrasco 2012). Indeed, the influential premotor theory of attention posits that covert orienting is nothing other than the preparation of an eye movement without its execution (premotor theory of attention, Rizzolatti et al. 1987; Sheliga et al. 1994; Sheliga et al. 1995).

In support of this tight coupling, closer inspection reveals that covert attention shifts may not be entirely covert. Although large scale eye movements may be absent during covert attention tasks, much smaller microsaccades can often be detected (for a review on the topic see: Hafed et al. 2021; Martinez-Conde et al. 2004), and have been suggested as a biomarker for covert attentional shifts (Engbert and Kliegl 2003; Hafed and Clark 2002; Laubrock et al. 2005; Pastukhov and Braun 2010; Tian et al. 2016; Tian et al. 2018). The precise nature of the coupling of covert and overt orienting processes is still disputed (for a review on the topic see: Belopolsky and Theeuwes 2012; Hunt et al. 2019; Klein and Pontefract 1994; Smith and Schenk 2012), but there are compelling similarities between the two ways of directing attention in space, which suggest that the underlying mechanisms might substantially overlap.

Given this theoretical context, we ask whether the phenomenon of SI, which arises with striking regularity in overt responses, extends to covert orienting. We operationalise covert orienting in terms of its perceptual consequences, specifically modulations in the ability to discriminate visual orientation at a cued or an uncued location (Fernández et al. 2019; Pestilli and Carrasco 2005). The key question was whether an irrelevant flash, which would induce SI in overt tasks, interferes with perceptual discrimination in a covert task. We adapted our previously-used double-step saccadic task (Buonocore et al. 2017c), to create a covert orienting task suitable for testing the inhibitory influence of an irrelevant visual transient. This effectively combined a classic task for the exogenous cueing of covert attention (Posner 1980) with a SI paradigm. We assume that the onset of the cue starts building-up activity at its location in space (i.e. covert orienting), which may subsequently be followed by reorienting to the opposite (uncued) location. Our initial hypotheses were based on the results of our earlier saccadic study (Buonocore et al. 2017c). We predict that the transient flash will interfere with covert orienting and that this effect will depend on the state of the attentional system, affecting performance only at specific CTOAs. If the pattern of covert orienting follows that of overt orienting, then we may expect this interruption to improve the ability to re-orient to the opposite (uncued) location, giving a relative enhancement of perceptual discrimination for invalidly cued targets at later CTOAs.

## Experiment 1 – Materials and Methods

### Participants

Fourteen participants (7 female), aged between 21 and 35 years (mean = 25.6, *SD* = 3.86), were included in the data analysis of Experiment 1. Four participants were excluded from the analysis based on poor perceptual performance (see Procedure). All participants were free from neurological and visual impairments. Experiment 1 was conducted with the approval of the University of Edinburgh Psychology Research Ethics Committee. Participants provided written, informed consent, and were financially compensated (at £10 per hour).

### Sample size considerations

SI in overt behavioural tasks is an extremely robust phenomenon, usually observed in every participant. Our previous studies of SI, like those of other researchers, have used relatively few participants (6-10), but many trials per condition (≥ 240) (e.g. Buonocore and McIntosh 2008; Buonocore and McIntosh 2012; Buonocore and McIntosh 2013; Buonocore et al. 2016; Buonocore et al. 2017c). These large trial numbers per condition are required for the estimation of SI parameters from a full distributional analysis of saccadic latencies; far fewer trials are necessary to estimate global performance measures (e.g. average latency).

The present covert experiment was designed as an exploratory follow up to our overt saccadic study, in which the addition of a flash was found to make saccadic reorienting more successful (Buonocore et al. 2017c). This effect was observed for all six participants in that study, with the flash increasing the probability of successful reorienting by an average of 10 percentage points (SD 3.4). This is a standardised effect size (dz) of 2.9, a huge effect statistically. Given uncertainty over how this effect would translate into orientation discrimination accuracy in a covert task in the present design, we did not perform a formal *a priori* analysis, but we aimed to at least double the sample size of our prior study. Eighteen participants were initially tested in Experiment 1, providing a final group of 14; no data analysis was conducted prior to termination of testing, except to determine participant exclusions.

### Experiment 1 – Apparatus and stimuli

Stimuli were generated using MATLAB R2017b (MathWorks, Inc.) and Psychophysics Toolbox 3.0.14 (Brainard 1997). All the stimuli were in grey scale on a grey background (20.6 cd/m^2^), presented on a 19-inch CRT monitor with a refresh rate of 75 Hz (13.3 ms temporal resolution). Participants were seated in front of the monitor at a viewing distance of 79.5 cm with their head on a chinrest and their eyes aligned with the centre of the screen. The fixation point was a small white dot of 0.05 degrees radius (84.9 cd/m^2^). The cue was a filled white circle of 0.2 degrees radius at full contrast (84.9 cd/m^2^), at an eccentricity of 10 degrees from the fixation point and 1.5 degrees above the centre of the possible probe location on that side (e.g. Pestilli and Carrasco 2005). The probe stimulus was a tilted Gabor patch with a radius of 0.55 degrees, a contrast of 0.2 and a spatial frequency of 1.78 c/deg, 10 degrees to the left or right of the fixation point. Eye movements were monitored with a tower-mount EyeLink 1000 system tracking the right eye at a sampling rate of 1000 Hz. Manual responses were recorded on a two-button response pad. The room was dark, except for the display monitor, and the operator monitor located behind the participant and facing away from them.

Each participant completed two testing sessions on separate days lasting about one hour each including breaks. The first testing session involved a practice block, a QUEST procedure for orientation threshold (repeated up to three times), and 38 blocks of experimental trials. The second testing session comprised 38 blocks of experimental trials.

### Experiment 1 – Procedure

At the beginning of each block, a 9-point calibration was conducted. Calibrations were repeated if the average error across all points was greater than 0.5 degrees, and after every 200 trials. In both the QUEST and the main experimental trials, participants were instructed to fixate the fixation point throughout the trial and to report the orientation of the Gabor patch (left or right tilted) by pressing the button under their left or right index finger respectively. We used discrimination of orientation, rather than spatial detection of the probe location, as a measure of change in sensitivity across the different conditions. In this way, the spatial propriety of the exogenous cue (location) was orthogonal to the features of the stimulus (orientation). Speed of responding was not emphasised. If participants moved their eyes outside of a fixation window of 3 degrees radius, the trial was aborted and randomly reshuffled into the remaining trial sequence.

In the first session, to familiarise with the basic task, participants first performed a practice block of 16 trials in which the onset of a cue was followed after a random delay by the onset of a probe at the same spatial location (congruent cue condition). The practice block was followed by a QUEST staircase block (Watson and Pelli 1983), a commonly used adaptive method for psychometric threshold. The QUEST block was used to identify a suitable orientation per participant that produced around 75% correct responses. Based on prior estimates of the psychometric function and the responses to different tilt intensities a final threshold was estimated.

The trial sequence for the QUEST followed the same structure as the subsequent experimental trials (description below), for a maximum of 80 trials, but only congruent cue conditions were used. The QUEST parameters were set to a 75% performance criterion, a beta of 1.5, and a grain of 1. If the estimated threshold orientation was greater than 15 degrees, the QUEST block was run again up to a maximum of three times. Participants were excluded from the main experiment if they still had an outcome greater than 15 degrees after the third QUEST block, or if their average performance in experimental trials was below 60% at the end of the whole first session. Four participants were excluded on this basis.

The experimental task was an adaptation of the classic cue Posner paradigm for covert perceptual judgements, modified to mimic the structure of Experiment 3 in Buonocore et al. (2017c) for overt eye movement responses (Fig. 1). Each trial began with the onset of a white fixation dot. After a random interval between 800 ms and 1200 ms, the cue was presented for 53.3 ms to the left or right of fixation. The probe was presented for 106.6 ms after a cue-target onset asynchrony (CTOA) selected on every trial from a uniform random distribution between 120 and 306.6 ms, with equal numbers of trials at the cued location (*congruent cue condition*) or at the uncued location (*incongruent cue condition*). On half of the trials, a black flash (0.34 cd/m^2^) covering the bottom and top thirds of the screen was presented for 13.3 ms simultaneously with the onset of the probe. This technique was introduced by Reingold and Stampe (2002) as a way to induce saccadic inhibition without interfering with the localization of a saccade target. It was recently adapted to saccade tasks requiring a concurrent perceptual response (Buonocore et al. 2016; Buonocore and Melcher 2015), establishing a lack of masking effects of this remote flash on probe perception.

**Figure 1.**
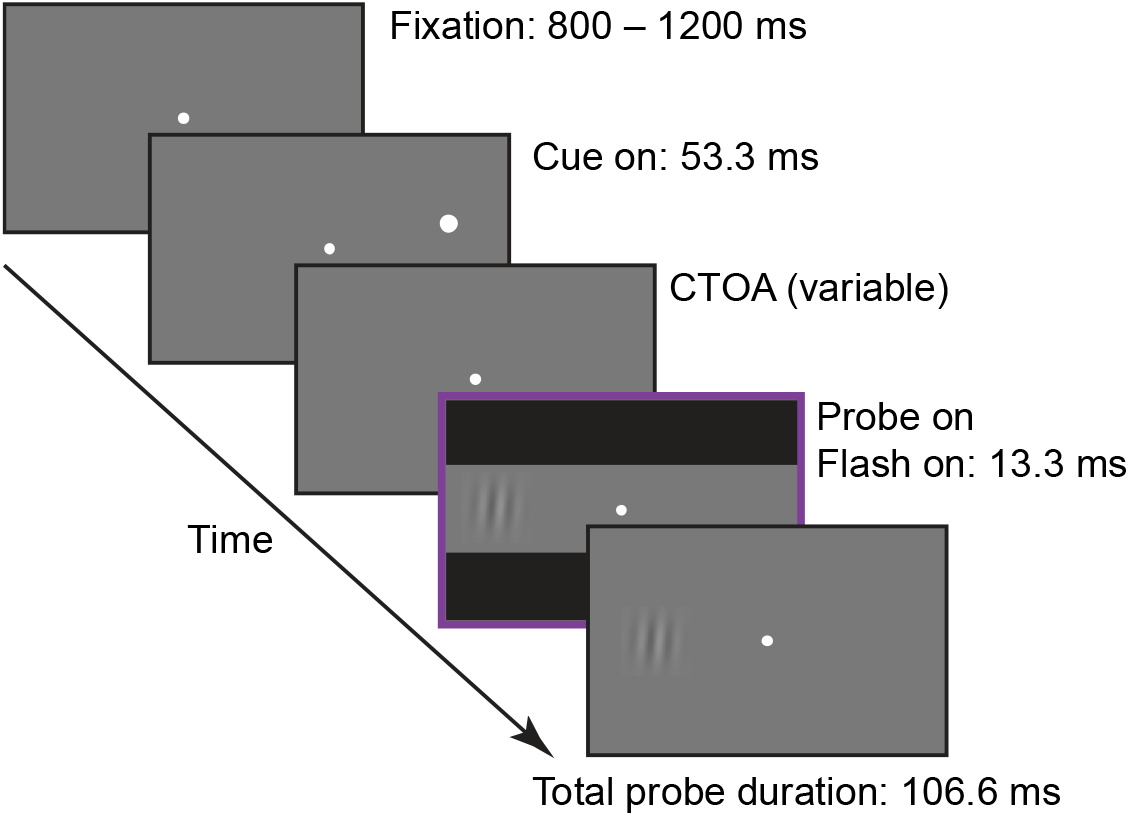
Trial scheme. Each trial started with the onset a fixation point. After a random interval between 800 to 1200 ms, a white dot cue was presented for 53.3 ms at 10 degrees to one side of fixation. After a random interval between 120 to 306.6 ms after cue onset (CTOA), the probe was presented for 106.6 ms either at the same (congruent) or the opposite (incongruent) location as the cue. On half of the trials, a flash appeared with the onset of the probe covering the bottom and top thirds of the screen for 13.3 ms. Here we show the example of an incongruent plus flash trial. Stimuli are not drawn to scale. Experiment 2 was identical to Experiment 1 but now in half of the trials the cue could be presented at the centre of the screen. We also used fixed CTOAs of 47, 82.3, 105.8 and 200 ms. The duration of the cue was slightly shorter than in Experiment 1, 35.2 ms, to allow an early CTOA of 47 ms. Flash duration was 11.3 ms and probe duration was 90 ms.

Participants completed 76 experimental blocks, for a total of 1216 valid trials, across the two test sessions. Within each block, there were 16 valid trials, created by the factorial combination of cue type (congruent, incongruent) by flash (no flash, flash) by probe side (left, right) by probe tilt (left, right). The first two factors (cue, flash) were of theoretical interest, whilst the latter two factors (probe side and tilt) were not, and were simply counterbalanced. The CTOA on each trial was drawn randomly from a uniform distribution between 120 and 306.6 ms. For data analysis, four CTOA categories were formed by grouping the CTOA in bins of 50 ms (centred at 140, 190, 240 and 290 ms). On average, each participant completed 76 trials per CTOA category for each combination of cue type (congruent, incongruent) by flash (no flash, flash).

### Experiment 1 – Data processing and analysis

Eye movements, saccades and microsaccades, were detected automatically based on velocity and acceleration thresholds of 15 deg/s and 450 deg/s^2^ respectively, then all trials were manually inspected and adjusted by an experienced researcher (AB). Samples in which the eye signal was unstable, lost or affected by blinks were flagged as “bad data” 2.5%. From the trials with good eye movement signals, we further excluded trials with manual reaction times less than 200 ms (4.1%) or more that 3.5 standard deviation above each participants average response latency (1.7%), leaving a total of 15682 good trials. To remove possible confounds in our results due to microsaccades (e.g. Hafed 2013), we also removed all trials with microsaccades detected between 200 ms before cue onset and 200 ms after probe onset (1604 trials removed, 10.22% of the good trials). This defined an interval of stable fixation of minimum duration 515 ms and maximum duration 715 ms, depending upon the CTOA. We checked if microsaccades were equally balanced in the congruent and incongruent cue condition and we found that on 47.5% microsaccades were executed in the congruent and 52.5% in the incongruent cue condition. Similarly, 48.1% microsaccades were detected in no flash trials and 51.9% in flash trials. For the remaining trials (N = 14078), incorrect responses were coded as 0 and correct responses as 1. We then used a mixed-effects logistic regression to test the influence of Cue type (congruent, incongruent), Flash (no flash, flash), and CTOA (four bins centred on 140, 190, 240, 290 ms) on perceptual performance.

To asses whether our stimuli were inducing inhibition, we also analysed microsaccade rates relative to both cue and probe onset. For each participant, we calculated for every trial the absolute frequency of microsaccades in bins of 20 ms within a time window spanning 200 ms before cue/probe onset and 500 ms after cue/probe onset. The absolute frequency was averaged across trials and converted to microsaccade rate by dividing it for the bin size (and transformed in seconds). Each microsaccade rate curve was then averaged across participants.

All data pre-processing and statistical analyses were conducted with custom scripts in MATLAB R2019a (MathWorks, Inc.) and R (R Core Team 2020) (lme4, lmerTest). The entire dataset is uploaded in the Open Science Framework archive at the following link: https://osf.io/9fnh4/?view_only=14b0ae67d49c4c1aa4898f66f20b473b

## Experiment 1 – Results

The present experiment follows up the recent findings in which we reported that inducing saccadic inhibition during oculomotor programming in a variant of a double-step paradigm (Becker and Jürgens 1979) could facilitate saccade reorienting behaviour (Buonocore et al. 2017c). Based on the findings for overt saccade responses, we asked if covert orienting, that we conceptualize in the framework of premotor preparation (Rizzolatti et al. 1987; Sheliga et al. 1994; Sheliga et al. 1995), could be subjected to similar inhibitory processes triggered by flash onset and which repercussions such inhibition of covert orienting would have on perceptual judgements. In our attentional version of the paradigm, cue onset, rather than accumulating build-up activity to trigger an eye movement, would initiate a similar build-up process that might increase visual sensitivity at its location. Usually this “shift in attention” is maximal after 100 ms relative to cue onset (e.g. Cheal and Lyon 1991) and it is followed by a reorienting of attention that decrease sensitivity at that location around 200 ms after cue (Cheal and Chastain 1999; Cheal et al. 1998; Posner and Cohen 1984; Pratt and Abrams 1999). Our expectation is that flash onset might hinder covert orienting capabilities in a specific time window that depends on the state of the attentional process, similar to how the sudden presentation of a visual stimulus can reset saccade programming only if it is presented at a critical time of the motor program, about 90 ms before its execution (Buonocore and McIntosh 2008; Reingold and Stampe 1999; Reingold and Stampe 2002). To test this idea, we ran a mixed-effects logistic regression to investigate how perceptual response accuracy (0,1) was modulated by *Cue* type (congruent, incongruent), *Flash* (no flash, flash), *CTOA* (140 ms, 190 ms, 240 ms, 290 ms, representing bin center) and their interactions. The model is summarized in Equation 1 in Wilkinson notation (Wilkinson and Rogers 1973):

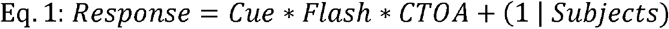

We found a strong main effect of *Cue* (*β* = -0.166, 95% CI = [-0.245, -0.086], t = -4.083, *p* = 0.001) confirming better orientation discrimination for congruently cued than for incongruently cued probes (Cameron et al. 2002). Overall, the cueing effect corresponded to an increase of about 3.5% in discrimination performance when cue location matched the probe location. For clarity, in Figure 2A we show the mean accuracy for each condition across all CTOAs. From the figure it is clear that on average the congruent cue conditions (green and blue lines) produced better orientation discrimination that the incongruent cue condition (yellow and purple lines). The analysis also revealed an effect of *Flash* (*β* = -0.1, 95% CI = [-0.177, -0.018], t = -2.406, *p* = 0.016) and an interaction between *Flash* and *CTOA* such that perceptual performance was disrupted in flash trials (Fig. 2A, blue and purple lines) compared to no flash ones (Fig. 2A, green and yellow lines), when the cue preceded the probe with a delay of 190 ms (*β* = -0.254, 95% CI = [-0.469, -0.039], t = - 2.314, *p* = 0.021) and 240 ms (*β* = -0.244, 95% CI = [-0.476, -0.012], t = -2.0636, *p* = 0.039). Figure 2B illustrates this interaction, plotting the difference between flash and no flash trials (irrespective of cue). Perceptual discrimination in the middle bins of the CTOA range was reduced by about 4%.

**Figure 2.**
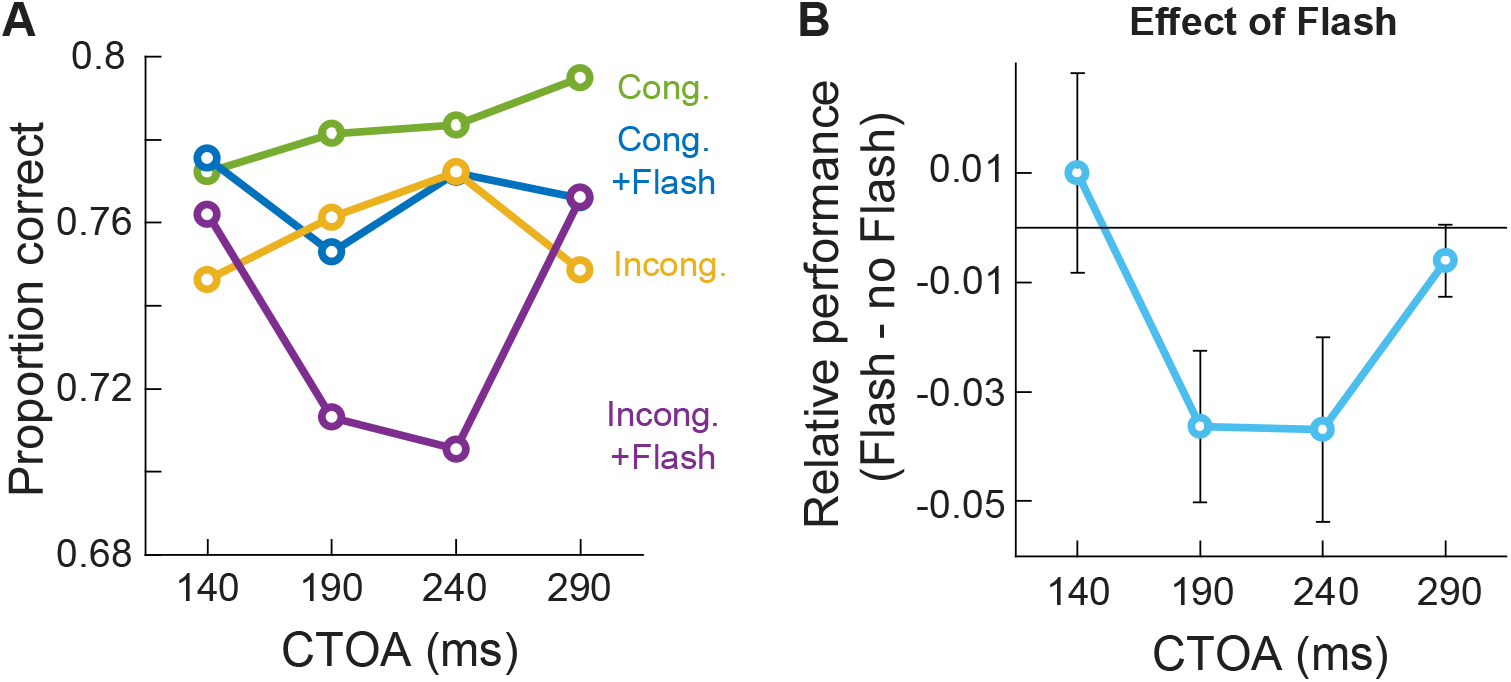
Perceptual performance in Experiment 1. **(A)** Congruent cues (green and blue lines) lead to better performance than incongruent cues (yellow and purple lines). Flash onset (blue and purple lines) interfered with orienting behaviour reducing accuracy relative to trials in which the flash was absent (green and yellow lines respectively). The effect was more pronounced when the cue was presented 190 to 240 ms before probe onset and for the incongruent condition (purple line). **(B)** Relative effect of the flash calculated as accuracy for flash trials minus no flash trials. A clear decrease in perceptual discrimination is visible for the middle range CTOAs. Error bars represent one standard error of the mean.

The data for each condition in Figure 2A suggests that the effect of the flash may be mostly driven by the incongruent cue condition (purple line), rather than the congruent cue condition (blue line). We explored this pattern by running two separate mixed-effects logistic regressions to test the influence of Flash and CTOA, and their interaction, on the congruent and incongruent trials. There were no significant effects for the congruent cue condition at any of the CTOAs, while there was an interaction in the incongruent cue condition with a strong effect of the flash at 190 ms (*β* = -0.331, 95% CI = [-0.632, -0.03], t = -2.155, *p* = 0.031) and 240 ms (*β* = -0.401, 95% CI = [-0.728, -0.074], t = -2.404, *p* = 0.016). While this analysis does not formally establish a three-way interaction, it is consistent with a stronger inhibitory influence of the flash in incongruent trials, relatively specific to the middle two time bins (190, 240 ms).

This pattern is evident in Figure 3, where we show the discrimination accuracies of our individual participants, contrasting the no flash against flash condition at each CTOA for the congruent (Fig.3A, top row panels) and the incongruent (Fig.3B, bottom row panels) cue conditions. The figure clarifies that perceptual performance in the congruent cue condition was not strongly affected at any CTOA. On the other hand, in the incongruent cue condition at the mid CTOAs there was a clear shift in perceptual performance (decrease in performance observed in 11 and 12 out of 14 participants respectively).

**Figure 3.**
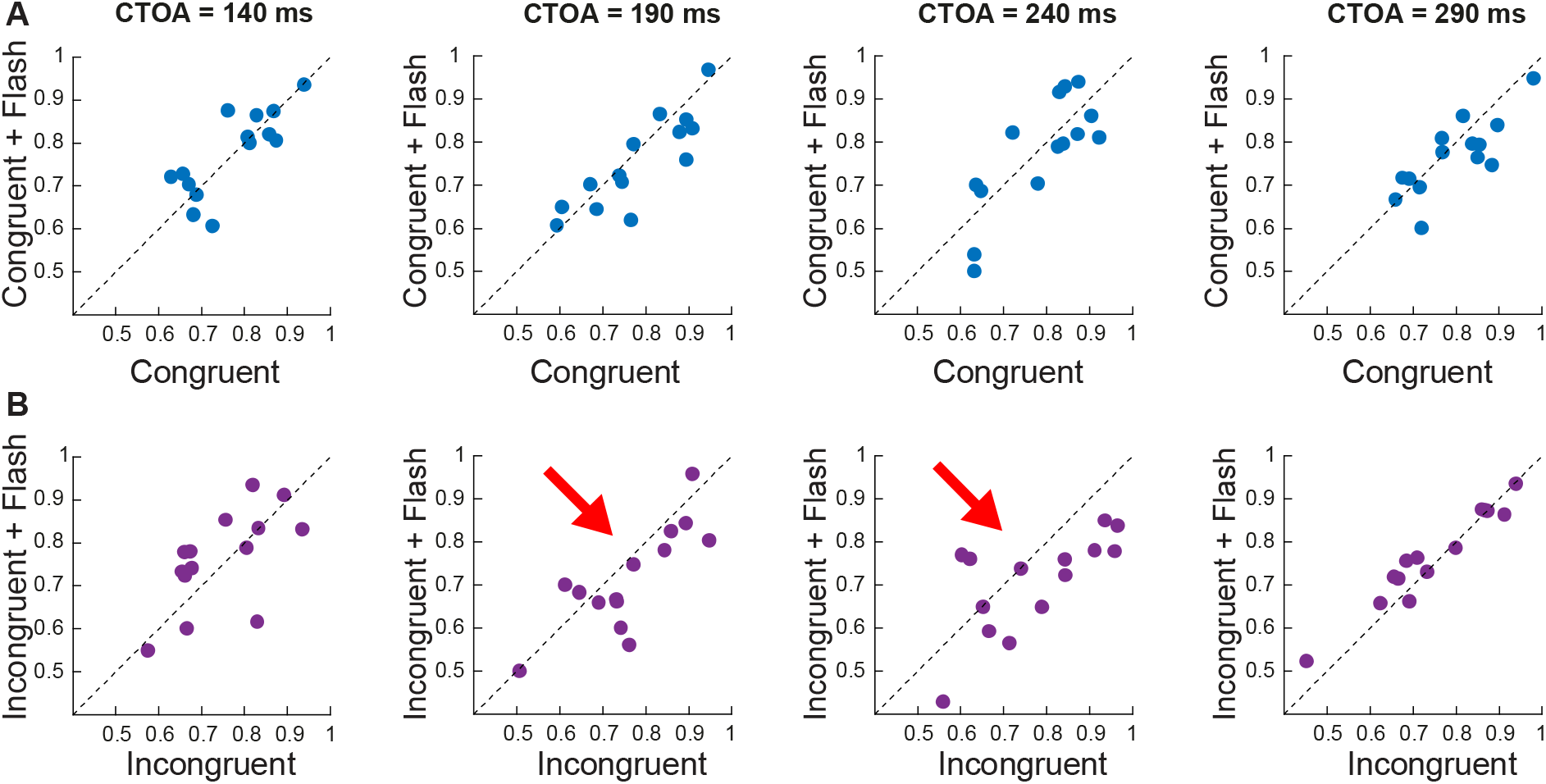
Raw data points of individual participants contrasting the no flash condition against flash condition at each CTOA for the congruent **(A)** and incongruent **(B)** cue condition. Dotted lines represents the unity slope line. At CTOAs of 190 ms and 240 ms (red arrow) there was a strong decrease in discrimination performance in the incongruent plus flash cue condition. Early and late CTOAs did not show any difference for trials with or without flash.

### Experiment 1 – Microsaccade analysis

In the previous section we showed that covert orienting was perturbed only when a flash was presented 190 to 240 ms after cue onset, in the total absence of eye movements. In the following analysis, we asked if the stimuli used in our paradigm were actually capable to inhibit overt responses and induce SI in a similar time period. To answer this question, we looked at microsaccades, the tiny eye movements (smaller than 1 degree) that continuously happen during covert attention tasks (Engbert and Kliegl 2003; Laubrock et al. 2005; Pastukhov and Braun 2010).

We detected a total of 3363 microsaccades in 2785 trials in a time period of 200 ms before cue onset to 500 ms after probe onset (1.2 microsaccades per trial within this time period). In Figure 4A, we aligned microsaccades to cue onset and, as expected, a strong microsaccadic inhibition took place between 200 to 400 ms after the cue (Buonocore et al. 2017a; Malevich et al. 2021; Rolfs et al. 2008; Tian et al. 2016; Tian et al. 2018), suggesting that the cue was strongly inhibiting overt orienting responses. We note that microsaccade rate was decreasing with a negative trend even before cue onset. This “microsaccade suppression” during the pre-cue interval indicates the effort of participants make to maintain fixation while waiting for the trial stimuli (e.g. Denison et al. 2019). In Figure 4B we aligned the same data to probe onset and calculated microsaccade rate separately for both no flash (green line) and flash (blue line) conditions. Overall, it is clear that microsaccadic inhibition induced by the cue was already so strong that microsaccades did not have time to recover by the time the probe was presented. Nonetheless, the inhibition was further enhanced 100 ms after probe onset and it was extended up to 300 ms thereafter (100 ms after the minimum reaction time allowed). Comparing microsaccadic rate in no flash (green) and flash (blue) conditions suggests that the flash did not have any additional measurable effect on the inhibition since it was already at ceiling. It is important to note that the lack of observable influence of the flash on overt behaviour is due to microsaccadic inhibition being total, but this does not mean that the flash should not have an influence on covert activation. Rather the fact that even the cue and probe were able to cause microsaccadic inbhibition implies that the much more salient flash should provoke even greater effects. Overall, the data confirm that our stimuli were indeed very powerful in inducing microsaccadic inhibition in a time period largely overlapping with the inhibition of covert orienting reported above.

**Figure 4.**
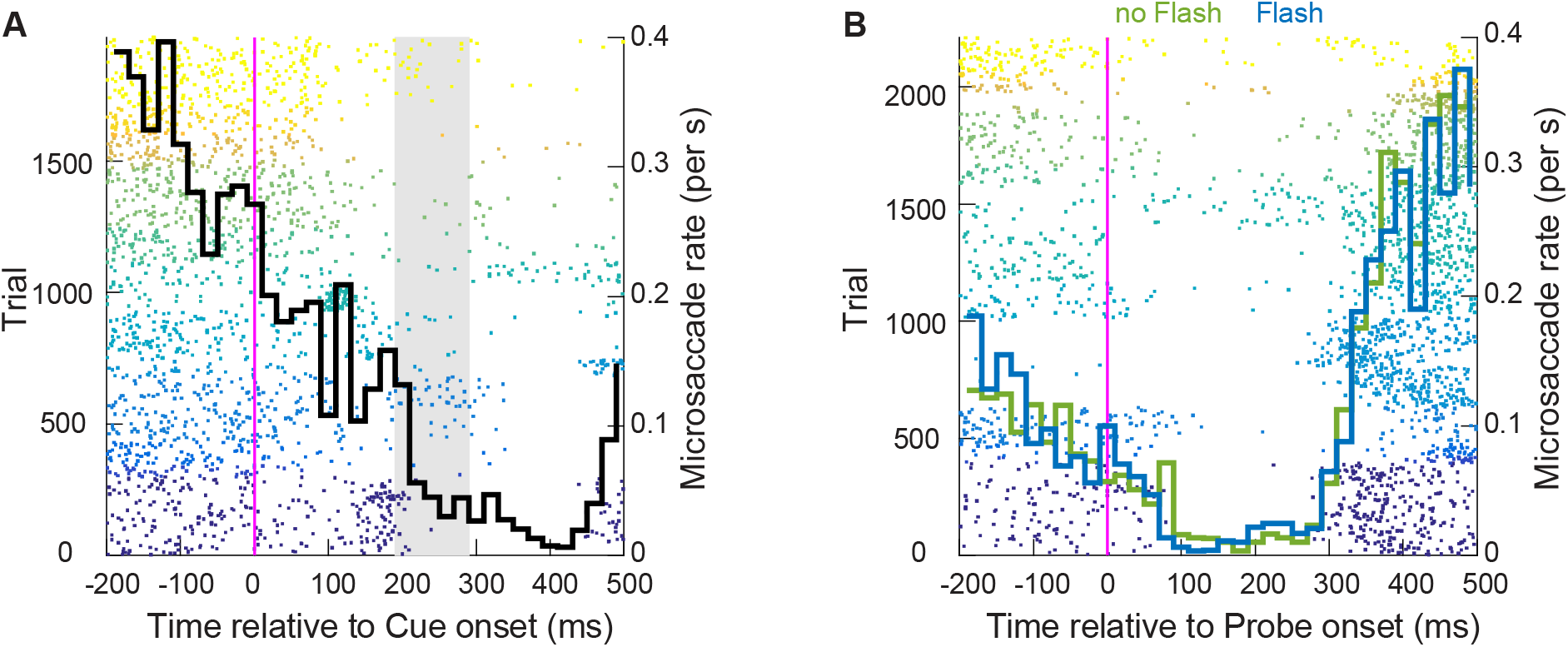
Microsaccadic inhibition in Experiment 1. **(A)** Microsaccade times aligned to cue onset. Each dot represents a microsaccade in an interval between 200 ms before to 500 ms after cue onset. Each colour band represents microsaccades for one participant. About 200 ms after cue onset, microsaccades are almost completely abolished. The black curve represents the averaged microsaccade rate across participants. The grey shaded area represents the time at which we recorded inhibition of covert orienting in absence of eye movements **(B)** Same as **(A)** but for microsaccades aligned to probe onset. Microsaccade rate is calculated separately for no flash (green line) and flash (blue line) conditions. Microsaccadic inhibition reached ceiling after the presentation of the probe alone, rendering difficult to see any clear additional effect of the flash.

## Experiment 1 – Interim discussion

The results of Experiment 1 suggest that covert orienting is inhibited by the flash, but only at certain time points, when the flash is presented between 190 to 240 ms after cue onset. This suggest that the only the reorienting component of the attentional shift, i.e. when attention is directed away from the previously cued location, was sensitive to the flash interference. In particular, we report that the effect was present only in the incongruent cue condition, when such reorienting process was exaggerated by the need to reorient attention to the opposite side of the visual field relative to the cue. The fact that we do not find any effect in the congruent cue condition at any of the CTOA suggests that the covert shift of attention was already completed by the time we presented the flash (e.g. Cheal and Lyon 1991) and attention was sufficiently allocated at the cue location rendering the discrimination process impervious to flash presentation. These results parallel the case in overt orienting in which a flash is presented at a time close to saccade onset, when the motor program for a saccade has passed a ‘point-of-no-return’, and is too advanced to be cancelled or delayed (Buonocore et al. 2016). One question that arises for the present covert case is whether the flash presented at the earliest CTOA (140 ms) completely missed the initial component of the orienting shift, not allowing any interference to take place, or if the early stage of covert orienting is somehow more resistant to inhibition.

To clarify at which stage of the attentional shift it is possible to inhibit orienting, we planned Experiment 2 where we tested the effect of the flash at the very early stages of the covert shift as well as at a later stage when reorienting is expected. Experiment 2 focused on the congruent cue condition only, by using a 100% informative cue (as opposed to 50% informative cues of Experiment 1). This design aim to mimic a condition in which the onset of a cue is immediately followed by a reflexive saccade. In the covert case, the cue acts as the stimulus to start attentional allocation to a particular point in space, similarly to a visual target starting the build-up activity for an eye movement. The subsequent flash aims to interfere with the ongoing premotor program. By using an early range of CTOAs compared to Experiment 1, we targeted an earlier window during the covert attentional shift.

## Experiment 2 – Material and Methods

Twelve participants (8 female), aged between 20 and 35 years old (mean = 25, *SD* = 4.2), were included in the data analysis of Experiment 2. Two participants were excluded from the analysis based on floor and ceiling effects (see *Experiment 2 – Procedure* below). All participants were free from neurological and visual impairments. Experiment 2 was conducted with the approval of relevant ethics committees at the Medical Faculty of Tuebingen University and of the University of Bielefeld. Subjects provided written, informed consent, and participated either as unpaid volunteers or for financial compensation (€10 per testing session).

### Sample size considerations

Experiment 2 was conducted during COVID restrictions in 2020-21, which made recruitment and testing opportunities extremely limited. We aimed to test equivalent participant numbers to Experiment 1, and managed to test 14 participants between two laboratories, though only 12 participants remained after exclusions. This was the maximum achievable sample size under the conditions that existed at the time.

### Experiment 2 – Apparatus and stimuli

Experiment 2 was similar to that for Experiment 1 but run in a different laboratory (Physiology of Active Vision Laboratory, Werner Reichardt Centre for Integrative Neuroscience, University of Tuebingen), with minor differences in setup^1^. The monitor was a 19-inch CRT monitor with a refresh rate of 85Hz (11.8 ms temporal resolution) positioned at a viewing distance of 57 cm. Stimuli were presented on a grey background (42.2 cd/m^2^). The fixation point was 0.01 degrees radius (97.3 cd/m^2^), and the cue (74.9 cd/m^2^) was presented either 1 degrees above the fixation point or at an eccentricity of 10 degrees from the fixation point and 1 degrees above the centre of the possible probe location on that side. All other stimulus parameters were identical to those in Experiment 1. Eye movements were monitored with a desktop mount EyeLink 1000 system, tracking the right eye at a sampling rate of 1000 Hz. Manual responses were recorded with two (left or right) of the five-button on a standard response pad (RESPONSEPixx, VPixx Technologies Inc.). Participants completed one testing session with one practice block and 16 blocks of experimental trials.

### Experiment 2 – Procedure

Experiment 2 procedure was similar to Experiment 1 except for a few critical modifications. After calibration, the experiment started with a practice block of 64 trials and no QUEST procedure was required. Instead, we used a probe stimulus with an orientation of either 84, 85, 86 degrees (right tilt) or 94, 95, 96 degrees (left tilt) chosen at random at the start of each trial. After a random interval between 800 ms and 1200 ms, the cue was presented for 35.2 ms either 10 degrees to the left or right of fixation or at the centre of the screen one degree above of the fixation point. The probe was presented for 90 ms after one of four randomly selected CTOAs: 47, 82, 106 and 200 ms, with equal numbers of trials at the cued location (*congruent cue condition*) or at the neutral location (*neutral cue condition*). The neutral cue condition was introduced as a control for any generalised temporal “warning effect” of the congruent cue (Coull and Nobre 1998; Hackley and Valle-Inclán 2003). On half of the trials, a black flash covering the bottom and top thirds of the screen for 11.3 ms was onset simultaneously with the probe.

Participants were asked to complete 16 blocks of experimental trials within the 75 minutes testing session. Within each block, there were 64 valid trials, one for each combination of cue type (congruent, incongruent) by flash (no flash, flash) by probe side (left, right) by CTOA (47, 82, 106 and 200 ms) by probe tilt (left, right). The first three factors (cue, flash, CTOA) were of theoretical interest, whilst the latter two factors (probe side and tilt) were not. Each participant completed 64 trials per CTOA for each combination of cue type (congruent, neutral) by flash (no flash, flash), except that some of the participants did not finish the last block due to time constrains (across subjects a total of 182 trials were not completed). Two of an original 14 participants were excluded from the analysis because their overall mean performance was either at floor (48.6%) or at ceiling (93.9%).

### Experiment 2 – Data processing and analysis

We collected a total of 12112 trials across all included participants in Experiment 2. We used identical criteria to Experiment 1 for trial exclusion. We rejected 1% of trials because of “bad data”, 0.17% trials with manual reaction times less than 200 ms and 1% of trials with manual reaction times more than 3.5 standard deviation above the participant’s average. We also remove 2.3% of trials in which critical stimulus flips were not properly timed with the monitor refresh rate. Our sample was of 11578 trials, from which we removed 1699 trials because of microsaccades (14.7%), leaving 9879 of microsaccade free trials in an interval between -200 ms before cue onset and 200 ms after probe onset (minimum saccade free interval equal to 250 ms and maximum saccade free interval equal to 400 ms). From the total of the excluded microsaccades, 50% belonged to the congruent and 50% in the incongruent cue condition. Similarly, 52.6% microsaccade were detected in no flash trials and 47.3% in flash trials. We ran a mixed-effects logistic regression to test the influence of Cue type (congruent, neutral), Flash (no flash, flash), and CTOA (47, 82, 105 and 200 ms) on perceptual performance. Paired-sample t-test were used to follow-up statistical analysis with an alpha level of 0.05. Microsaccade analysis was identical to Experiment 1.

All data pre-processing and statistical analyses follow the procedure described in Experiment 1 and the entire dataset is uploaded in the Open Science Framework (see *Experiment 1 - Data processing and analysis*).

## Experiment 2 – Results

In Experiment 2 we tested whether the inhibitory effect of the flash could occur within the range relevant to early orienting or if it was only effective in the later (reorienting) range. To do so, we tested orientation discrimination for probes presented at congruent cue locations in a time window between 50 to 200 ms. To control for possible effects driven only by the temporal aspects of the cue, rather than its location in space, we also tested a neutral cue condition in which a spatially non-informative cue was presented centrally. Based on the results of Experiment 1, we expected to see modulations of discrimination performance at the time of reorienting, when attention moves away from the cued location, and/or at early CTOAs, when facilitation is expected. To explore these patterns, we ran a mixed-effects logistic regression with *Cue* type (congruent, neutral), *Flash* (no flash, flash), *CTOA* (47 ms, 82 ms, 106 ms, 200 ms), and their interactions (same model as Eq. 1).

The first significant effect was an interaction between *Flash* and *CTOA* (Fig. 5A). In flash trials there was an enhancement of about 4% in discrimination performance when the probe was presented 47 ms after the cue compared to later CTOAs (CTOA 82 ms: *β* = - 0.036, 95% CI = [-0.32, 0.247], t = -0.252, *p* = 0.801 ; CTOA 106 ms: *β* = -0.287, 95% CI = [-0.566,-0.008], t = -2.017, *p* = 0.044; COTA 200 ms: *β* = -0.242, 95% CI = [-0.528, 0.042], t = -1.67, *p* = 0.095). Since the interaction was irrespective of cue type, i.e. congruent or neutral, it suggests that the flash was having a general “warning effect”, increasing visual sensitivity when presented immediately after the cue (Coull and Nobre 1998; Hackley and Valle-Inclán 2003).

**Figure 5.**
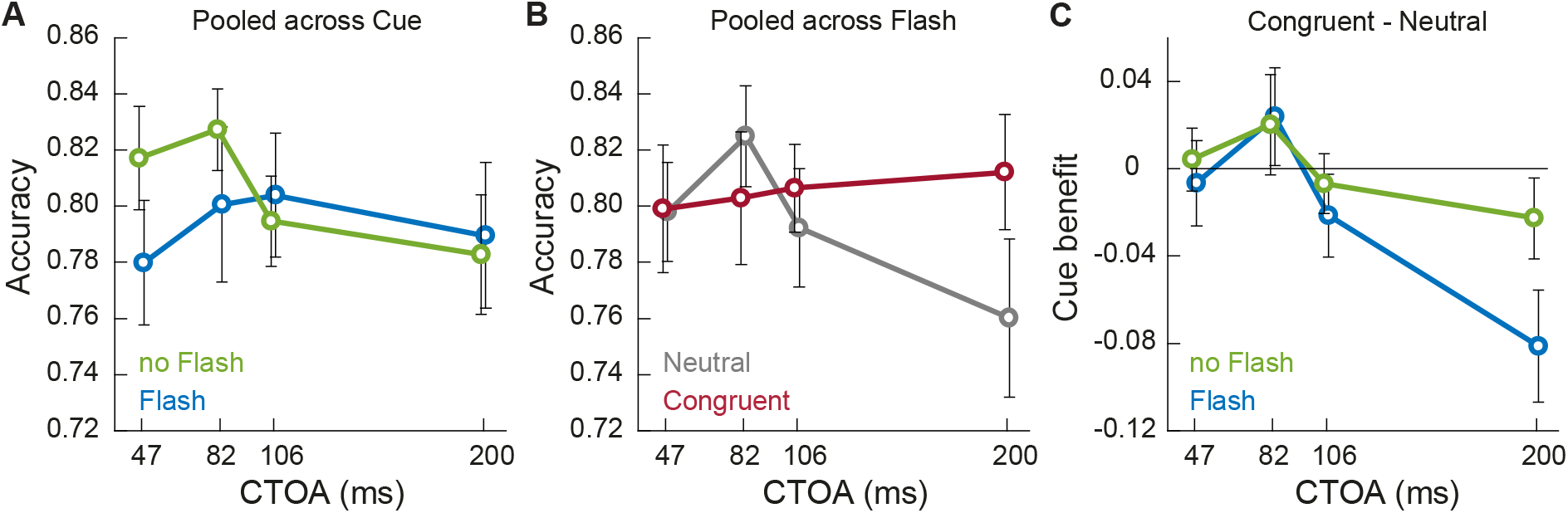
Perceptual performance. **(A)** At the early CTOA of 47 ms, performance is boosted by the presence of the flash (blue lines) compared to no flash trials (green lines) and irrespective of the cue type (**B)** Neutral cues (grey lines) do not modulate perceptual performance across CTOAs. On the other hand, congruent cues (red line) have an impact on discrimination performance for the late CTOA of 200 ms, where visual sensitivity for the incoming probe is impaired. **(C)** At the time of reorienting, i.e. 200 ms, the presentation of the flash (blue line) affects perceptual discrimination capabilities by impeding reorienting and maintaining visual sensitivity higher compared to no flash trials (green line).

We also found an interaction between *Cue* and *CTOA* (Fig. 5B). While performance in the neutral cue condition was stable at about 80% (grey line), the congruent cue condition (red line) was modulated across CTOAs with a decrease in discrimination performance of about 4% taking place 200 ms after cue onset ms (*β* = 0.298, 95% CI = [0.013, 0.583], t = 2.049, *p* = 0.041). Numerically, perceptual performance was slightly better at CTOA 82 ms, but overall we did not see a significant facilitation effect at any of the early CTOAs. This result suggested that in our paradigm the attentional shift triggered by the cue at early CTOAs was not strong enough to boost performance significantly for this task. On the other hand, attentional allocation started reducing at cue location at around 200 ms after cue onset, compatible with previous reports (Cheal and Chastain 1999; Cheal et al. 1998; Posner and Cohen 1984; Pratt and Abrams 1999) and our results from Experiment 1.

The significant decrease in discrimination performance observed in the congruent cue condition at 200 ms after cue onset (Fig. 5B) was suggestive of attention moving away from the cue location (reorienting). This result motivated us to explore in more detail if the flash could interfere with this part of the orienting process, as observed for the incongruent cue condition in Experiment 1. To do so, we computed a measure of cue-benefit by subtracting accuracy in the neutral cue condition from that in the congruent cue condition in both no flash and flash trials at each CTOA. Paired-sample t-test showed that only at 200 ms was there a difference between flash and no flash conditions (t(11) = -2.94, p=0.013), with discrimination performance being about 6% higher in flash trials (Fig. 5C). This result supports the hypothesis that the flash has a temporal window of action in which it can inhibit attention moving away from cue location, similarly to the result reported in Experiment 1.

### Experiment 2 – Microsaccade analysis

We detected a total of 3454 microsaccades over 3133 trials in a time period of 200 ms before cue onset to 500 ms after probe onset (1.1 microsaccades per trial within this time period). Similarly to Experiment 1, microsaccade rate (black line) was completely disrupted about 100 ms after cue onset (Fig. 6A). When data were aligned to probe onset (Fig. 6B) and microsaccade rate was split by flash condition (green: no flash, blue: flash), despite microsaccadic rate being close to zero for both conditions, there is indication of a slightly larger inhibition for flash trials. As in Experiment 1, microsaccadic inhibition after the probe was long lasting, with the rebound period starting only at around 300 ms after probe onset. This results confirms that our stimuli could abolish any subsequent eye movement in a time period compatible with the covert effect recorded in absence of eye movement.

**Figure 6.**
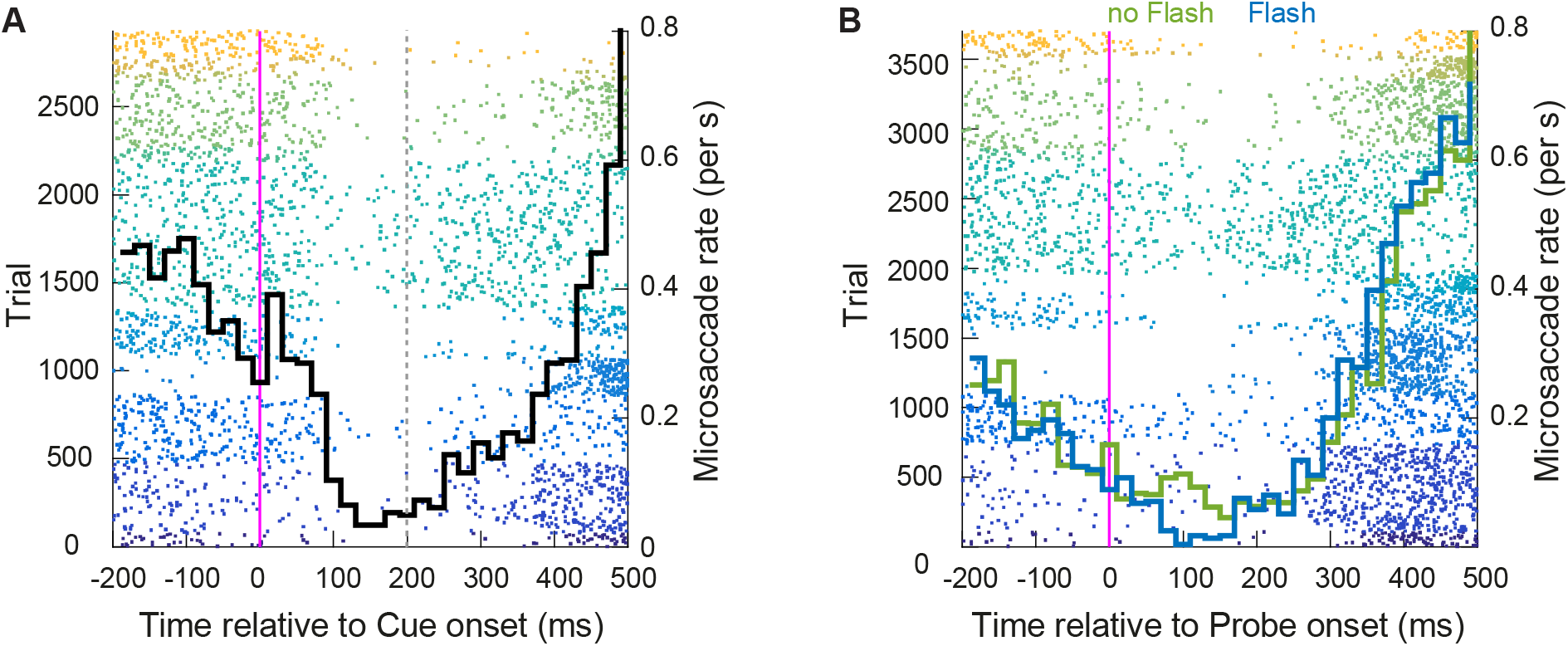
Microsaccadic inhibition. Same colour conventions as Figure 5. **(A)** After cue onset, microsaccades are almost completely abolished (black line). The vertical grey line indicates the CTOA were we recorded the inhibition of covert reorienting. **(B)** Same as **(A)** but for microsaccades aligned to probe onset. There is a slightly stronger inhibition in flash (blue line) compared to no flash (green line) trials.

## Experiment 2 – Interim discussion

The results from Experiment 2 confirm that perceptual discrimination can be impaired by the flash during the reorienting stage of the attentional shift. At the early CTOAs we did not observe any specific effect of the flash in the congruent cue condition. This result is surprising because within this time range covert orienting should mostly overlap with the preparation of an eye movement to a lateralised cue, similarly to what happens in pre-saccadic shifts of attention (Buonocore et al. 2017b; Deubel and Schneider 1996; Hoffman and Subramaniam 1995; Kowler et al. 1995; Li et al. 2016; Li et al. 2019; Ohl et al. 2017; Remington 1980; Rolfs and Carrasco 2012; Shepherd et al. 1986). By presenting a flash at this stage of motor programming should be possible to strongly interfere with the build-up activity at the cued location, as we clearly see in SI paradigms. One possible explanation is that the early cue was not powerful enough to induce strong orienting toward the cue location, as indicated by the absence of a facilitation effect for early cues. On the contrary, we see a general facilitation effect in flash trials for CTOA equal to 47 ms. The fact that we see such facilitation speaks instead of a temporal “warning effect” of the flash, independent of cue type, that increased visual sensitivity for the incoming probe (Coull and Nobre 1998; Hackley and Valle-Inclán 2003). This facilitation might have overshadowed any inhibitory signal associated with flash presentation.

The effect recorded at CTOA equal 200 ms in the congruent cue condition might be the result of the flash introducing a small delay in the process of attentional reorienting. Consequently, a higher level of visual sensitivity was maintained at the cued location for a longer time, increasing perceptual performance. This result is consistent with the flash interfering with the incongruent cue conditions of Experiment 1. In this case, visual sensitivity was maintained higher at the incongruent cue location, leading to a decrease in performance for a probe presented at the opposite cue side. Taken together, Experiment 1 and 2 suggest that the later reorienting component can be effectively perturbed by a generalised flash, while the early orienting component of the attentional shift may be less susceptible to inhibition.

## General conclusions

In the present manuscript, we uncovered a new phenomenon within a classic cueing paradigm (Cameron et al. 2002; Posner 1980) in which the visual discrimination of a probe stimulus was deteriorated by the simultaneous presentation of a brief flash event, but only when the flash was preceded by the exogenous cue between 190 to 240 ms. In Experiment 1, the effect was particularly emphasised for the condition in which cue location was incongruent with the location of the incoming probe impeding reorienting toward the probe and decreasing perceptual discrimination. In Experiment 2, a similar inhibitory effect was found in a congruent cue condition, by inhibiting reorienting away from the cue and maintaining performance slightly higher. Taken together, we suggest that the build-up of attentional resources at a cue location has a specific temporal window in which an inhibitory signal can interfere, leading to an inhibition of covert reorienting. However, the exact time course and aspects of the covert orienting process subject to interference require further investigation.

We assume that the mechanisms behind the inhibitory process might share similar characteristics with the well-known phenomenon of SI for overt responses in which a flash (or other transient event) interferes with the preparation of any eye movement that is due about 90 ms later (Bompas and Sumner 2011; Buonocore and McIntosh 2008; Buonocore and McIntosh 2012; Edelman and Xu 2009; Reingold and Stampe 1999; Reingold and Stampe 2002). Despite the differences in the time course of SI and the inhibition of covert orienting reported here, we suggest that the two phenomena might rely on the same neural circuitries, in accordance with the mechanistic overlap between covert and overt orienting proposed by the premotor theory of attention (Rizzolatti et al. 1987; Sheliga et al. 1994; Sheliga et al. 1995). From a neuronal perspective, the onset of the flash might decrease the firing rate of the premotor build-up SC neurons coding for saccades to a particular location in space (Buonocore et al. 2021; Dorris et al. 2007; Munoz and Istvan 1998) and consequently introduce a small delay in the premotor plan. The source of the inhibitory signal might be an increased in activation of OPNs following flash onset (Buonocore et al. 2020; Buonocore et al. 2017a), which is then broadcast to the SC and other upstream visual and visuo-motor areas. Higher order visual areas tuned for stimulus features such as orientation (e.g. V1, Hubel and Wiesel 1959) might then be affected by the temporal delays introduced in the early premotor process for target selection, affecting perceptual discrimination.

One might ask why if the neural mechanisms are shared between covert and overt processing there are temporal differences between the two inhibitory effects. We suggest that perhaps overt SI has an intrinsic “all-or-nothing” nature, because a saccade either is or is not launched within a specific time window. Any depression of the premotor activation can prevent the saccade from launching at the expected time, leading to a large SI effect 90 ms after flash onset. On the other hand, covert orienting might depend on a more gradual readout of premotor activity, which transiently changes following flash onset but continues to accumulate until a discrimination is made. For this covert process to provide sufficient cumulative activation for discrimination could require a longer integration period compared to the process to trigger saccade execution. Moreover, while SI can be almost entirely modelled within a small network within subcortical oculomotor areas such as SC and the lower brainstem, visual discrimination is a process that involves higher order cortical visual area that contribute to reshape the temporal profile of the effect. This account is speculative, but the more general argument is that superficial differences in timing do not necessarily preclude an underlying identity of the covert and overt inhibition phenomena, because the measured consequences may be strongly shaped by the very different readout processes associated with saccade execution and perceptual discrimination.

Based on the general SI framework (Reingold and Stampe 2002) and our previous findings on saccades (Buonocore et al. 2017c), we expected flash onset to interfere with covert orienting behaviour in a specific time window, inducing an SI-like inhibition effect. Beyond this, if the pattern of covert orienting was strictly following that of overt orienting as observed in our previous study, we would have expected this interruption to improve the ability to re-orient to the opposite (uncued) location, giving a relative enhancement of perceptual discrimination for incongruently cued targets. In this respect, the effect would have been the perceptual counterpart to the higher rate of successful reorienting saccades observed when SI was boosted in “flash-jump” trials in the overt task. The other prediction was that perceptual performance at congruent cue locations might have been slightly impaired. Although our results confirm that the effect of the flash depended upon the stage of evolution of the covert shift, the direction of the effect was contrary to our initial expectation. Rather than impairing early orienting behaviour, the flash impaired later reorienting. This finding can perhaps be informed by the idea that reorienting following interruption after a transient event carries a small temporal cost (see reorienting latency for Experiment 3, Table 3 in: Buonocore et al. 2017c). In our covert paradigm the flash might have introduced a similar delay during covert orienting, requiring more time to redirect to the probe location. For example, in Experiment 1, given the brevity of the probe stimulus to be discriminated (106 ms), this small delay may have been enough to reduce resources at the probe location and consequently deteriorate sensitivity for the probe stimulus (Salinas and Stanford 2018). That is, while orienting (to the cued location) in our design might have been mostly or wholly completed within 100 ms (e.g. Cheal and Lyon 1991), and so invulnerable to interruption by a flash, reorienting (in incongruent trials) occurred later, and was vulnerable to interruption, reducing the opportunity to process the brief probe stimulus. This mechanism can also accommodate the outcome of Experiment 2, where the flash inhibited reorienting in the congruent cue condition. In this case, the orienting component of the shift was also completed by 100 ms and attentional allocation might have started moving away from the cued location. Introducing a small interruption in this reorienting process by presenting a brief flash might have maintained more resources at the cued location, increasing perceptual performance.

It is important to emphasise that the interruption effect was restricted to later CTOA times (190-240 ms in Experiment 1 and 200 ms in Experiment 2), despite the fact that the flash was always simultaneous with the probe. This rules out the possibility that passive masking mechanisms (Alpern 1952; Breitmeyer and Ogmen 2000) could account for the deterioration of probe discrimination; if masking were responsible, then the appearance of the flash simultaneous with the probe would always lead to the same impairment, across all experimental conditions. Moreover, in Experiment 2 we found that the flash presented at early CTOAs enhanced performance, acting as a general “warning” signal for the incoming probe and increasing the overall sensitivity of the visual system. This result is also incompatible with possible passive masking effect and speaks in favour of the flash interference being specific for the attentional state.

Taking together these observations, we can draw some predictions from our data to apply to future research work on inhibition of covert orienting. One main hypothesis is that the flash would always alter perceptual performance when it occurs during a sensitive period of the covert process triggered by the cue. The sensitive period could be predicted by estimating the time at which the covert shift would engage or disengage from the cue. This feature is in fact similar to the time-locked interference of the flash relative to saccadic reaction times. Finally, inhibition of covert orienting is then expected to also extend to paradigms in which pre-saccadic shift of attention are involved, potentially altering the strong benefits of coupling covert orienting with eye movements. We suggest that this paradigm may lead to new insights into how the attentional shift develops over time, and help to investigate the extent to which covert and overt processes share underlying neural processes.

## Footnotes

1 Five out of the fourteen participants were collected at the University of Bielefeld with the setup matched as closely as possible, but with a different monitor refresh rate (100 Hz).

